# Biological insights from plasma proteomics of non-small cell lung cancer patients treated with immunotherapy

**DOI:** 10.1101/2024.01.02.573876

**Authors:** Jair Bar, Raya Leibowitz, Niels Reinmuth, Astrid Ammendola, Eyal Jacob, Mor Moskovitz, Adva Levy-Barda, Michal Lotem, Rivka Katzenelson, Abed Agbarya, Mahmoud Abu-Amna, Maya Gottfried, Tatiana Harkovsky, Ido Wolf, Ella Tepper, Gil Loewenthal, Ben Yellin, Yehuda Brody, Nili Dahan, Maya Yanko, Coren Lahav, Michal Harel, Shani Raveh Shoval, Yehonatan Elon, Itamar Sela, Adam P. Dicker, Yuval Shaked

**Affiliations:** Institute of Oncology, Chaim Sheba Medical Center, Tel Hashomer, Israel; and Faculty of Medicine, Tel Aviv University, Tel-Aviv, Israel; Shamir Medical Center, Oncology Institute, Zerifin, 70300, Israel; and Faculty of Medicine, Tel Aviv University, Tel Aviv, Israel; Asklepios Kliniken GmbH, Asklepios Fachkliniken Muenchen, Gauting, 82131, Germany; and the German Center for Lung Research (DZL); OncoHost LTD. Binyamina, Israel; Thoracic Cancer Service, Rabin Medical Center Davidoff Cancer Centre, Beilinson Campus, Petah Tikva, 4941492, Israel; Biobank, Department of Pathology, Rabin Medical Center - Beilinson Campus, Petah Tikva, 4941492, Israel; Center for Melanoma and Cancer Immunotherapy, Hadassah Hebrew University Medical Center, Sharett Institute of Oncology, Jerusalem, 9112001, Israel; Kaplan Medical Center, Rehovot, 7642002, Israel; Institute of Oncology, Bnai Zion Medical Center, Haifa, 3339419, Israel; Oncology & Hematology Division, Cancer Center, Emek Medical Center, Afula, 1834111, Israel; Department of Oncology, Meir Medical Center, Kfar-Saba, 4428164, Israel; Barzilai Medical Center, Faculty of Health Sciences, Ben-Gurion University of the Negev, Ashkelon, Israel; Division of Oncology, Tel-Aviv Sourasky Medical Center, Tel Aviv, 6423906, Israel; Department of Oncology, Assuta Hospital, Tel Aviv, 6971028, Israel; Department of Radiation Oncology, Thomas Jefferson University, Philadelphia, Pennsylvania, United States of America; Faculty of Medicine, Technion – Israel Institute of Technology, Haifa, Israel

## Abstract

**Introduction:** Immune checkpoint inhibitors have made a paradigm shift in the treatment of non-small cell lung cancer (NSCLC). However, clinical response varies widely and robust predictive biomarkers for patient stratification are lacking. Here, we characterize early on-treatment proteomic changes in blood plasma to gain a better understanding of treatment response and resistance.

**Methods:** Pre-treatment (T0) and on-treatment (T1) plasma samples were collected from 225 NSCLC patients receiving PD-1/PD-L1 inhibitor-based regimens. Plasma was profiled using aptamer-based technology to quantify approximately 7000 plasma proteins per sample. Proteins displaying significant fold changes (T1:T0) were analyzed further to identify associations with clinical outcomes. Bioinformatic analyses of upregulated proteins were performed to determine potential cell origins and enriched biological processes.

**Results:** The levels of 142 proteins were significantly increased in the plasma of NSCLC patients following ICI-based treatments. Soluble PD-1 exhibited the highest increase, with a positive correlation to tumor PD-L1 status. Bioinformatic analysis of the ICI monotherapy dataset revealed a set of 30 upregulated proteins that formed a single, highly interconnected network with CD8A serving as a central hub, suggesting T cell activation during ICI treatment. Notably, the T cell-related network was detected regardless of clinical benefit. Lastly, circulating proteins of alveolar origin were identified as potential biomarkers of limited clinical benefit, possibly due to a link with cellular stress and lung damage.

**Conclusions:** Our study provides insights into the biological processes activated during ICI-based therapy, highlighting the potential of plasma proteomics to identify mechanisms of therapy resistance and potential biomarkers for outcome.

## INTRODUCTION

Over the past decade, immune checkpoint inhibitors (ICIs) directed against the programmed cell death protein-1 (PD-1) and programmed death ligand-1 (PD-L1) axis have emerged as the standard of care in the treatment of non-small cell lung cancer (NSCLC) and other advanced malignancies (1). These agents disrupt the interaction between PD-1 expressed on T cells and PD-L1 found on tumor cells, leading to an enhanced antitumor immune response (2). ICIs are considered a breakthrough for the management of NSCLC, primarily due to their ability to elicit durable responses, and in some cases amounting to cure of metastatic disease. The use of ICIs in the early-disease, pre-operative setting clearly decreases recurrence rates and increases cure rates (3). However, despite the impressive long-term survival observed in selected ICI-treated patients, the overall response rate remains modest, with only 20-30% of NSCLC patients experiencing durable benefit (4) and the rest displaying intrinsic or acquired resistance to ICIs (5). Resistance mechanisms include aberrations in cellular signaling pathways, the exclusion of T cells from the tumor microenvironment, the infiltration of immunosuppressive cell populations, the presence of inhibitory checkpoints, dampened interferon-γ signaling, histological transformations, and loss of tumor-associated antigenic proteins (6–8). However, these mechanisms are hard to detect, highlighting a need for clinical predictive biomarkers of response and resistance to therapy.

Various tissue-based biomarkers, including PD-L1 expression in tumor cells, high tumor mutational burden, and lymphocytic tumor infiltrates, have been explored as predictive biomarkers for immunotherapy response. However, clinical evidence demonstrates moderate predictive performance for these biomarkers (9, 10). There is intense interest in blood-based biomarkers due to their minimally invasive nature and potential for allowing longitudinal monitoring (11). For example, several studies demonstrate the value of monitoring circulating cell-free tumor DNA (ctDNA) during treatment to predict clinical response to ICI-based regimens in NSCLC patients (12–14). With respect to baseline blood-based biomarkers, low baseline levels of soluble PD-L1 (sPD-L1) have been linked to improved objective response rates in ICI-treated NSCLC, melanoma, and renal cell carcinoma patients (15, 16), and an elevated absolute neutrophil-to-lymphocyte ratio at baseline has been associated with poorer survival outcomes in NSCLC patients receiving anti-PD-1 therapy (17). Nonetheless, as individual entities, these biomarkers may not exhibit a consistently high degree of predictive accuracy as they do not reflect the multi-faceted nature of response and resistance to ICIs. Blood plasma proteomic profiling is a powerful approach for biomarker discovery with the potential to capture the heterogenous mechanisms of response and resistance to therapy (18, 19), and aid the development of personalized treatment strategies based on ICIs in combination with another agent. Additionally, the dynamic nature of the plasma proteome allows for temporal monitoring of immune system activity and disease progression (20). Thus, plasma proteomic profiles may provide real-time insight into tumor-immune system dynamics, thereby serving as a rich source of potential predictive biomarkers.

Our previous studies elucidated how host-mediated responses to various cancer treatment modalities contribute significantly to disease progression and the development of therapy resistance (21). Most recently, utilizing preclinical murine models, we demonstrated that ICI agents induce systemic alterations in host-derived factors, subsequently bolstering tumor aggressiveness, with a pivotal role attributed to interleukin (IL)-6 in this cascade (22). In a previous clinical study of ICI-treated NSCLC patients, we analyzed nearly 800 plasma proteins and identified CXCL10 and CXCL8 as predictive biomarkers for treatment response along with clinical parameters (23). Despite its limited cohort and protein dataset size, our clinical study underscores the potential of plasma proteomic profiling to stratify patients based on therapeutic response.

In the present study, we used an aptamer-based assay to measure approximately 7000 plasma proteins in pre- and on-treatment plasma samples from NSCLC patients treated with ICI-based regimens, allowing for the characterization of proteomic changes upon treatment. The analysis revealed: (i) ICI-induced elevation in soluble PD-1 (sPD-1) in the circulation; (ii) a unique plasma proteomic signature associated with T cell activation; and (iii) circulating proteins possibly originating from alveolar cells as potential blood-based biomarkers for limited therapeutic benefit from ICIs. Overall, our study provides mechanistic insights into therapy resistance, paving the way towards biomarker discovery.

## MATERIALS AND METHODS

### Sample collection

Blood plasma samples and clinical data were collected from advanced-stage NSCLC patients as part of a clinical study (PROPHETIC; NCT04056247). All clinical sites (Asklepios Kliniken GmbH, DE; Thoraxklinik at University Hospital Heidelberg, DE; Hadassah Medical Center, IL; Tel Aviv Sourasky Medical Center, IL; Rambam Health Care Campus, IL; Bnai Zion Medical Center, IL; Meir Medical Center, IL; Sheba Medical Center, IL; HaEmek Medical Center, IL; Kaplan Hospital, IL; Barzilai Medical Center, IL; Rabin Medical Center, IL; Assuta Medical Centers, IL; and Shamir Medical Center, IL) received IRB approval for the study protocol and all patients provided written informed consent. Patient blood samples were drawn at baseline (referred to as T0) and, on average, 4 weeks after the treatment commenced, prior to the second dose of treatment (referred to as T1). Blood samples were drawn into tubes containing EDTA as an anticoagulant, and plasma was separated from the whole blood by centrifuging at 1200 x g at room temperature for 10-20 minutes within 4 hours of venipuncture. This protocol allows for a maximum of 4 hours between sample collection and plasma separation to accommodate the multi-center nature of the clinical trial. The plasma supernatant was stored at -80°C and later shipped frozen to the analysis lab. The protocol adheres to the Clinical and Laboratory Standards Institute (CLSI) guidelines and aligns with the SomaLogic plasma collection guidelines, similar to (24). Clinical parameters including age, ECOG, sex, line of treatment; treatment type; Tumor Proportion Score (TPS); histological type; clinical benefit at 3 months, and overall survival (OS) was obtained for all patients. Patients were classified as ‘responders’ (R) or ‘non-responders’ (NR) based on whether they achieved clinical benefit at 3 months (according to RECIST 1.1). Thus, the R population included patients displaying complete response, partial response or stable disease and the NR population included patients with progressive disease.

### Proteomic profiling

Plasma samples from 225 patients were analyzed using the SomaScan® V4.1 assay that measures 7596 protein targets, of which 7288 are human proteins. The SomaScan technology uses slow-off-rate modified DNA aptamers that bind target proteins with high specificity to quantify the concentration of proteins in a sample. Results are provided in relative fluorescence units (RFU). The analysis was conducted at OncoHost’s CLIA-certified laboratory in North Carolina and SomaLogic’s laboratory in Colorado.

### Statistical and bioinformatic analysis

Data were analyzed using Python, R, and Cytoscape. The plasma levels per protein were expressed as fold change values (T1:T0 ratio) on a log2 scale. Differences in fold changes were assessed using paired t-tests with Benjamini-Hochberg FDR correction (FDR<0.01 for comparisons in the entire cohort, as well as in ICI monotherapy or combination therapy sub-cohorts, and FDR<0.1 for comparisons between R and NR patients in the ICI monotherapy sub-cohort). Pathway enrichment analysis was performed with gProfilier, WebGestalt and ShinyGo 0.77 web tools (25–27) on relevant sets of proteins (i.e., proteins displaying significant fold changes within the entire patient cohort or patient subcohorts) with Benjamini-Hochberg FDR <0.05 against the background of approximately 7000 proteins, using a cut-off as indicated in the figures. ShinyGo 0.77 was used to generate networks of enriched pathways (showing the hierarchical relationship between enriched pathways) where two pathways were connected if they shared at least 20% of proteins. VolcaNoseR was used to generate and explore the data with volcano plots (28). Protein-protein interaction (PPI) networks were generated using the STRING version 12.0 database with a 0.9 confidence interaction score (29).

Multivariate analysis was used to identify associations between sPD-1 fold change and categorical clinical data (age, ECOG, sex, line of treatment; treatment type; Tumor Proportion Score (TPS); histological type; clinical benefit at 3 months, and OS). Cox proportional hazards models were used to analyze the association between the protein of interest and OS.

## RESULTS

### Patient characteristics

The study included 225 NSCLC patients treated with ICI-based therapy. One hundred patients received PD-1/PD-L1 inhibitor monotherapy and 125 received a PD-1/PD-L1 inhibitor combined with chemotherapy (**Table S1**). Patient demographics and clinical parameters are presented in **Table 1**. The mean age was 66.4 years. Approximately a third of the patients were female. The majority (69.3%) of patients had adenocarcinoma and 20.4% of the patients had squamous cell carcinoma. Overall, 39.6%, 23.6% and 26.7% of patients had tumors with PD-L1 expression of ≥50%, 1-49% and <1%, respectively. Clinical benefit at 3 months was evaluated according to RECIST criteria where patients with complete response, partial response, and stable disease were termed responders (R), and those with progressive disease were termed non-responders (NR). Based on this definition, the cohort comprised 78.4% R and 21.5% NR patients. The high proportion of R patients in our cohort can be explained by the early time point at which clinical benefit was assessed (i.e., at 3 months) and the inclusion of stable disease in the R population in this study. To allow comparisons between different patient populations, the study included an additional cohort of NSCLC patients treated with chemotherapy alone (n=20) and a cohort of melanoma patients treated with ICI-based therapy (n=20). Patient characteristics of these cohorts are shown in **Tables S2 and S3**.

**Table 1:**
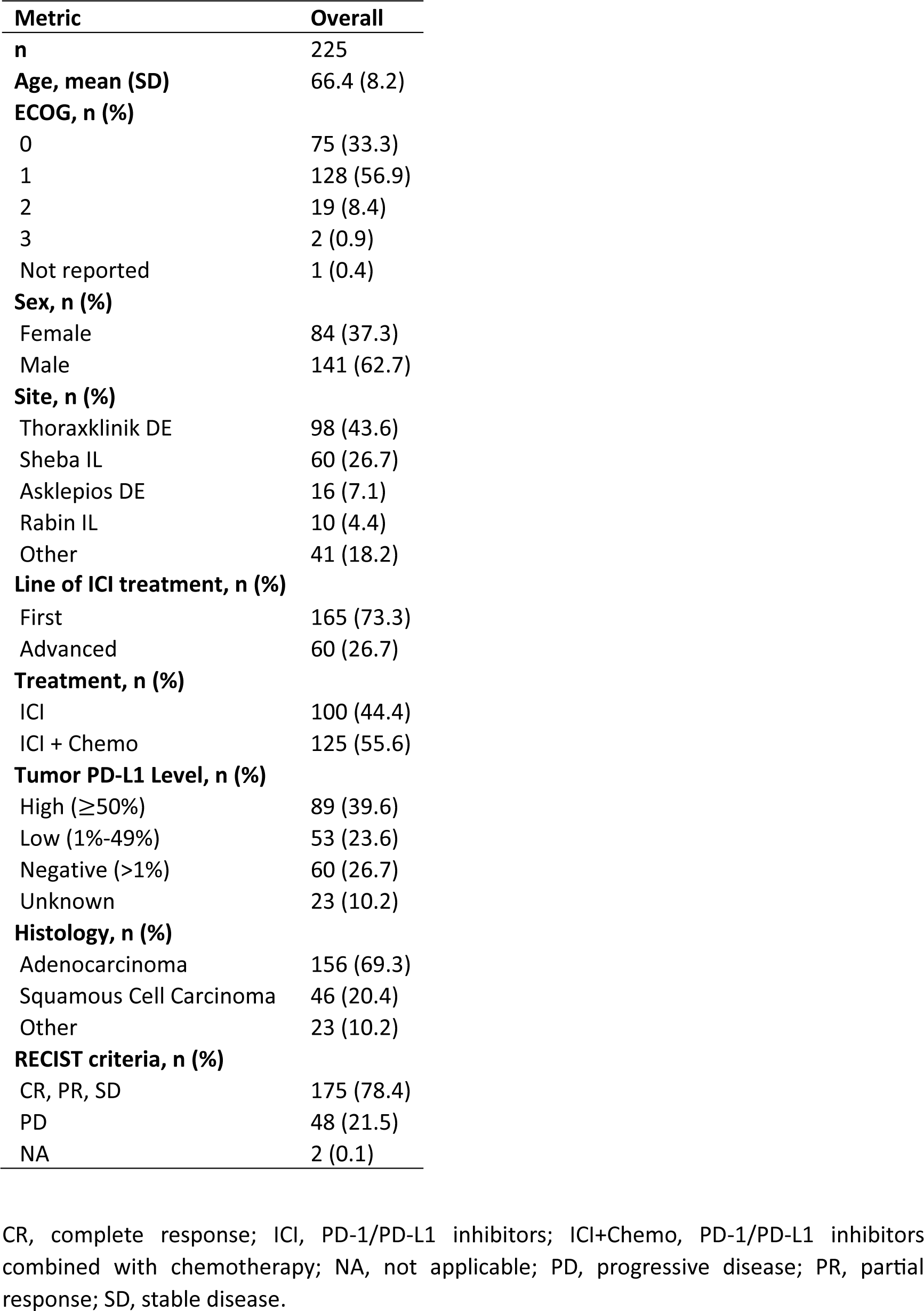
Patient demographics, clinicopathological characteristics and outcomes.

| **Metric** | **Overall** |
| --- | --- |
| **n** | 225 |
| **Age, mean (SD)** | 66.4 (8.2) |
| **ECOG, n (%)** |  |
| 0 | 75 (33.3) |
| 1 | 128 (56.9) |
| 2 | 19 (8.4) |
| 3 | 2 (0.9) |
| Not reported | 1 (0.4) |
| **Sex, n (%)** |  |
| Female | 84 (37.3) |
| Male | 141 (62.7) |
| **Site, n (%)** |  |
| Thoraxklinik DE | 98 (43.6) |
| Sheba IL | 60 (26.7) |
| Asklepios DE | 16 (7.1) |
| Rabin IL | 10 (4.4) |
| Other | 41 (18.2) |
| **Line of ICI treatment, n (%)** |  |
| First | 165 (73.3) |
| Advanced | 60 (26.7) |
| **Treatment, n (%)** |  |
| ICI | 100 (44.4) |
| ICI + Chemo | 125 (55.6) |
| **Tumor PD-L1 Level, n (%)** |  |
| High ($\geq 50\%$) | 89 (39.6) |
| Low (1%-49%) | 53 (23.6) |
| Negative ($>1\%$) | 60 (26.7) |
| Unknown | 23 (10.2) |
| **Histology, n (%)** |  |
| Adenocarcinoma | 156 (69.3) |
| Squamous Cell Carcinoma | 46 (20.4) |
| Other | 23 (10.2) |
| **RECIST criteria, n (%)** |  |
| CR, PR, SD | 175 (78.4) |
| PD | 48 (21.5) |
| NA | 2 (0.1) |

CR, complete response; ICI, PD-1/PD-L1 inhibitors; ICI+Chemo, PD-1/PD-L1 inhibitors combined with chemotherapy; NA, not applicable; PD, progressive disease; PR, partial response; SD, stable disease.

### ICI therapy elevates systemic levels of soluble PD-1

To study changes in circulating proteins during treatment with ICI-based therapy, pre-treatment (T0) and on-treatment (T1) patient plasma samples were profiled using the Somalogic aptamer-based assay that measures ∼7000 proteins per sample. The proteomic data were expressed as fold-change values of T1 to T0 measurements. The plasma levels of 142 proteins (represented by 164 aptamers) were significantly increased upon treatment (Log2FC>0.1 and FDR q value <0.01). Most notably, sPD-1 was highly elevated in plasma upon treatment, as demonstrated by measurements from the two aptamers that detect this protein (FDR q value 6.04E-88 and 3.66E-61, **Figure 1A-B**). The aptamers, designated 15623-1 and 9227-15, bind to regions located within aa 1-167 and aa 27-167 of sPD-1, respectively, corresponding to regions encoded by exon 1 and 2 in the PD-1 gene. Thus, both aptamers recognize the full-length membrane-bound protein and the soluble forms created by proteolytic cleavage or alternative splicing (personal communication with Somalogic). Analysis of patient subgroups revealed that sPD-1 fold change was highest in patients receiving PD-1 inhibitor therapy with or without chemotherapy, where the monotherapy subgroup displayed a significantly higher fold change than the combination therapy subgroup. In contrast, no fold changes were found in patients receiving PD-L1 inhibitors alone, PD-L1 inhibitors combined with chemotherapy, or chemotherapy alone (**Figure 1C**). Since ICI therapy targets the PD-1-PD-L1 axis, we next asked whether plasma levels of sPD-L1 are affected by ICI-based therapies. sPD-L1 plasma levels remained unchanged upon PD-1 inhibitor monotherapy, PD-1 inhibitors with chemotherapy, and chemotherapy alone, whereas they dropped significantly upon PD-L1 inhibitor therapy with or without chemotherapy (**Figure 1D**). This drop may be explained by the binding of the therapeutic anti-PD-L1 antibody to the aptamer binding site within the sPD-L1 molecule which may limit its detection by the assay or promote its clearance by immune mechanisms. This effect presumably does not occur in the case of anti-PD-1 therapeutic antibodies and sPD-1 molecules. Lastly, a correlation was found between sPD-1 fold change in plasma and baseline PD-L1 expression in tumors. Specifically, in patients receiving ICI monotherapy (i.e., PD-1 or PD-L1 inhibitors as single agents), fold-change of sPD-1 was significantly higher in the PD-L1 ≥50% group in comparison to the PD-L1 <1% group, whereas no differences between the PD-L1 groups were observed in patients receiving a combination of ICI with chemotherapy (**Figure 1E**). Overall, our findings demonstrate that sPD-1 plasma levels are elevated upon treatment with PD-1 inhibitors but not PD-L1 inhibitors. The addition of chemotherapy to ICI regimens dampens the on-treatment elevation in sPD-1 plasma levels, possibly due to the immunosuppression effect of chemotherapy on immune cells, among them T cells (30).

**Figure 1:**
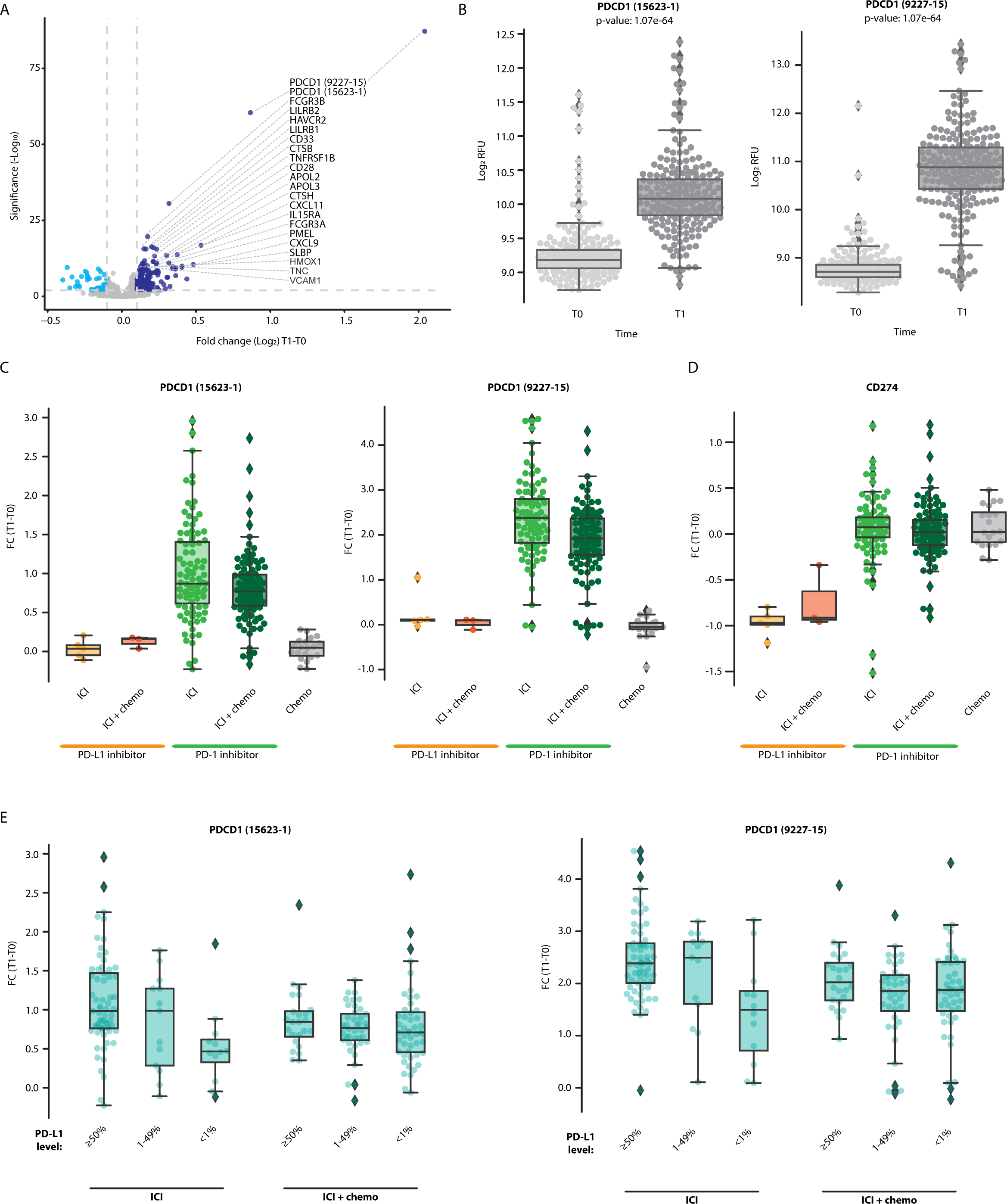
sPD-1 is elevated in the plasma following treatment with ICI-based therapy and is associated with tumor PD-L1 expression. A. Volcano plot representing fold changes (T1:T0) in plasma protein levels in patients treated with ICI-based therapy (n=225). Proteins displaying significant positive (dark blue) and negative (light blue) fold-change are indicated. B. Plasma levels of sPD-1 at T0 and T1, as quantified by two sPD-1 aptamers (15623-1 and 9227-15). C-D. Fold change of sPD-1 (C) and sPD-L1 (D) in patients receiving PD-1/PD-L1 inhibitor monotherapy (ICI), PD-1/PD-L1 inhibitors in combination with chemotherapy (ICI+chemo), or chemotherapy alone. E. Patients were grouped according to baseline tumor PD-L1 expression levels (≥50%, 1-49%, <1%). The fold changes of sPD-1 in patients treated with PD-1/PD-L1 inhibitor monotherapy (ICI) and PD-1/PD-L1 inhibitors in combination with chemotherapy (ICI+chemo) are shown.

### On-treatment elevation in sPD-1 plasma level is associated with improved overall survival

We next investigated whether sPD-1 fold change is associated with various patient demographic or clinical parameters. No significant associations were found between sPD-1 fold change and patient age, line of treatment or clinical benefit at 3 months (**Table S4**). To explore possible associations further, we compared sPD-1 fold change in R and NR patients within ICI monotherapy and ICI-chemotherapy groups. In the ICI monotherapy group, sPD-1 fold change was slightly higher in the R population than in the NR population, although the difference was not statistically significant. No differences were found between R and NR populations within the ICI-chemotherapy group (**Figure 2A**). To assess whether sPD-1 fold change is associated with long-term outcomes, we used the median sPD-1 fold change as a threshold to classify patients as having high or low sPD-1 fold change and compared overall survival (OS) between the two patient groups. In ICI monotherapy cases, patients with high sPD-1 fold change displayed longer OS than patients with low sPD-1 fold change, indicating its predictive power for survival rather than for early clinical benefit (**Figure 2B**). In contrast, no difference in OS was observed in patients receiving combination ICI-chemotherapy (**Figure 2C**). Lastly, no differences in OS were found between patients grouped by sPD-L1 fold-change (**Figure S1**), suggesting that sPD-1 is a superior indicator for improved OS compared to sPD-L1, at least in our experimental set-up.

**Figure 2:**
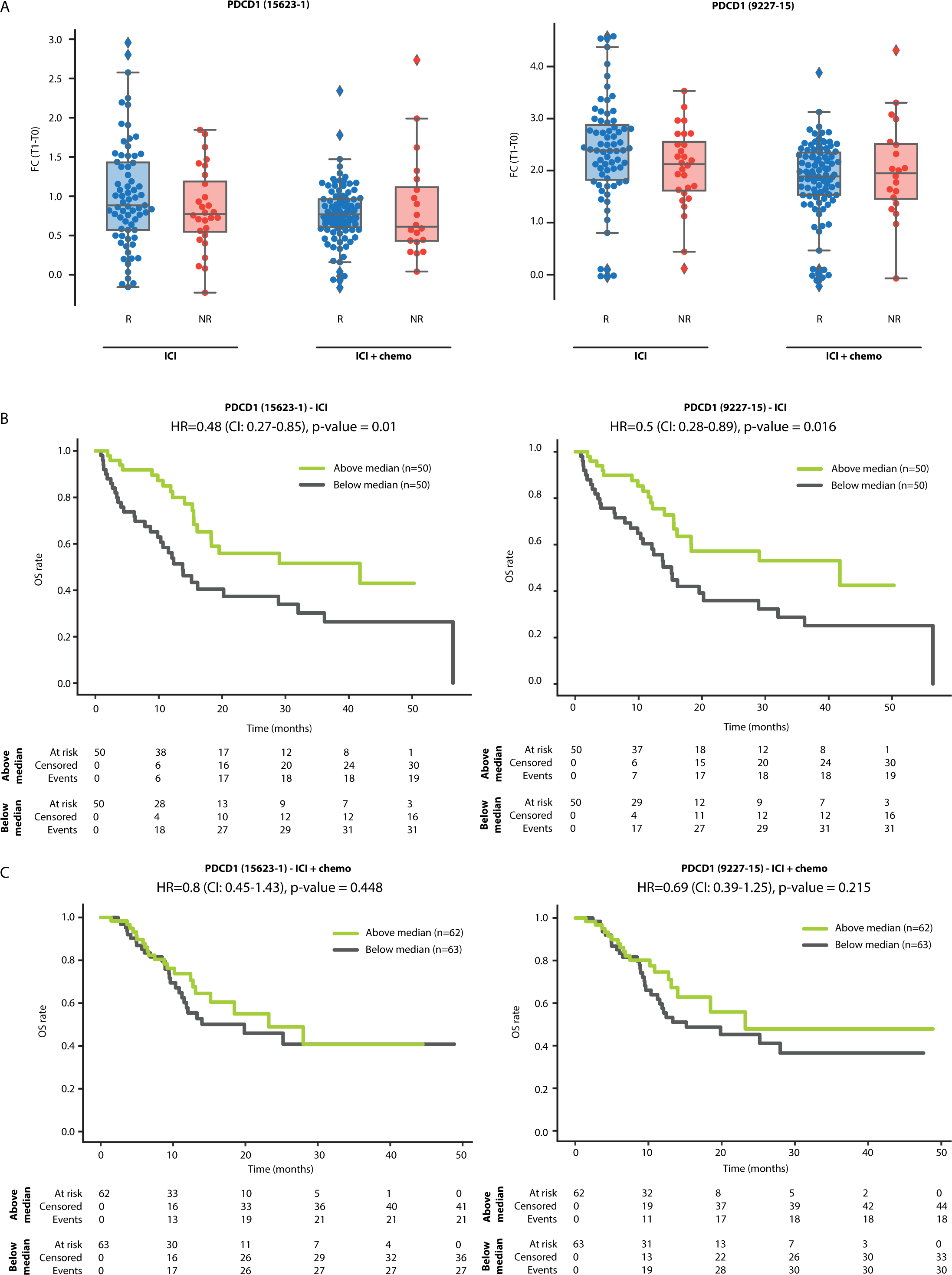
sPD-1 fold change is associated with overall survival. A. Patients were classified as responders (R) or non-responders (NR) according to clinical benefit at 3 months. Shown are fold changes (T1:T0) of sPD-1 in R and NR patients treated with PD-1/PD-L1 inhibitor monotherapy (ICI) and PD-1/PD-L1 inhibitors in combination with chemotherapy (ICI+chemo). B-C. Kaplan-Meier plots showing the relationship between sPD-1 fold change and overall survival (OS) in patients treated with PD-1/PD-L1 inhibitor monotherapy (ICI; B) and PD-1/PD-L1 inhibitors in combination with chemotherapy (ICI+chemo; C). Patients were classified as having high or low sPD-1 fold change using the median sPD-1 fold change as the classification threshold.

### ICI therapy elevates plasma levels of proteins associated with T cell activity irrespective of clinical benefit

To gain insight into the biological processes induced by PD-1/PD-L1 inhibitors, we first analyzed proteomic data from NSCLC patients receiving ICI monotherapy, to avoid the potential disruptive effect of chemotherapy. Within this dataset, the plasma levels of 85 proteins (represented by 100 aptamers) were found to be significantly increased upon treatment (log2FC>0.1 and FDR<0.01, **Figure 3A**). Pathway enrichment analysis of biological processes associated with these proteins revealed T-cell activation, proliferation, and differentiation (**Figure 3B-C**, and **Table S5**). Specifically, among the 85 proteins, 30 were associated with a strong T cell network with a confidence level of 0.9 (**Figure 3D**). Among these 30 proteins are the known T cell immune checkpoint proteins, LAG3, CD28, PD1 and HAVCR2, as well as other immune checkpoints, including LILRB2, and TREM2 (31). In addition, IL2R, IL15R, and Granzyme A, were also enriched in this T cell network, further indicating T cell activation and proliferation (32). Likewise, increased plasma levels of B2M and HLA-C suggest T cell activation through antigen presentation (33, 34). Notably, most proteins are cell-secreted and plasma membrane proteins (**Figure S2**). Interestingly, the T cell network was detected in both R and NR populations, albeit by different sets of proteins, indicating that at least based on circulating factors, T cell-related biological processes occur regardless of the therapeutic outcome (**Figure S3**). We found that the T cell network was completely disrupted in the patient subgroup receiving combination ICI-chemotherapy (**Figure S4**), further indicating that adding chemotherapy to ICI regimens weakens ICI-induced T cell-related processes. We further compared the proteins displaying positive fold changes upon ICI monotherapy and ICI-chemotherapy regimens. T cell-related processes were highly enriched in the set of proteins found exclusively in the ICI monotherapy group. In contrast, no enrichment was detected in the collection of proteins found solely in the ICI-chemotherapy group. The 27 proteins in both groups yielded an intermediate enrichment, suggesting that T cell-related processes are primarily associated with ICI treatment (**Figure S5**). Overall, on-treatment changes in the plasma proteome suggest a robust T cell-related biological process in both R and NR populations. This effect is prominent in patients receiving ICI monotherapy whereas it is not detected in patients receiving combination ICI-chemotherapy.

**Figure 3:**
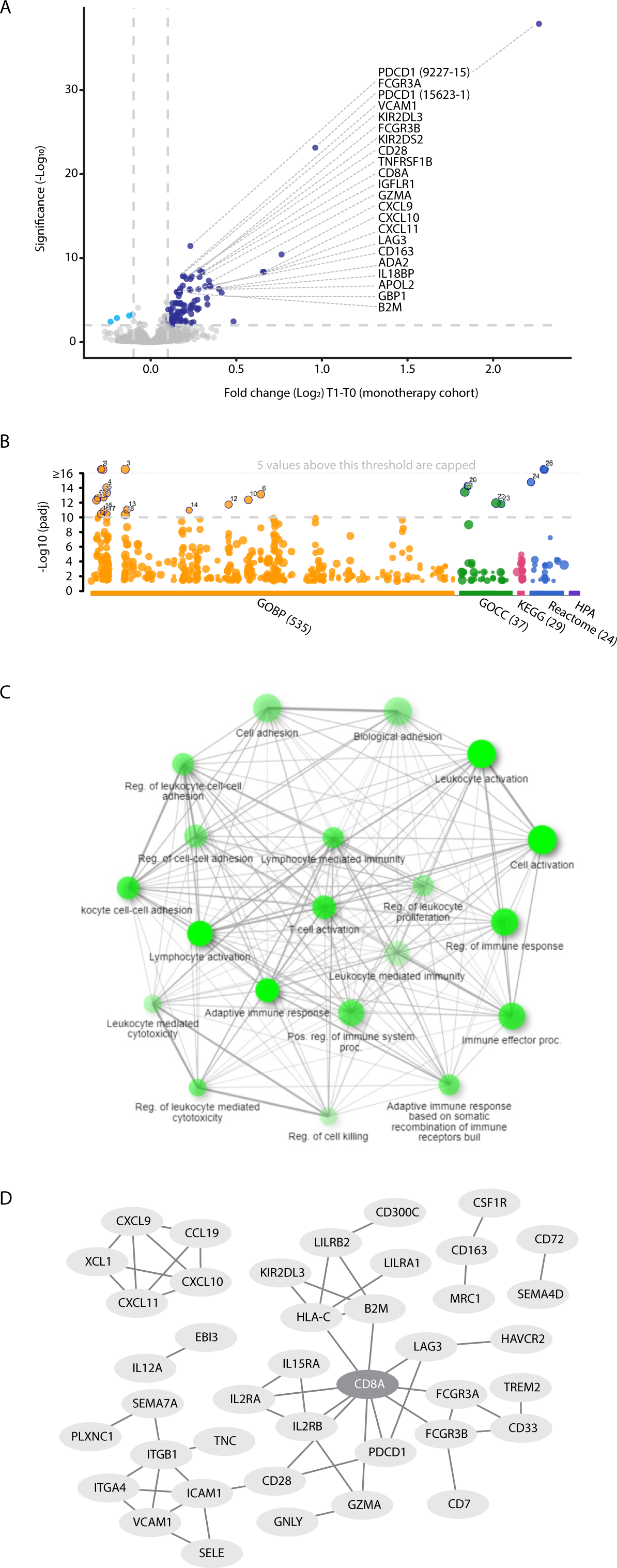
Proteins associated with T cell-related processes are elevated in the plasma during ICI treatment. A. Volcano plot representing fold change (T1:T0) in plasma protein levels in patients treated with PD-1/PD-L1 inhibitor monotherapy (n=100). B. Pathway enrichment analysis of proteins exhibiting significant and positive fold change. Categorization is based on gProfilier. Cut-off was set to -log10(q value) =10. C. Network of GO biological processes showing relationships between enriched pathways using ShinyGo. Notably, T cell-associated pathways are highly represented. D. The protein-protein interaction network shows the T cell hub (interaction score with 0.9 confidence).

### Elevated plasma levels of intracellular proteins correlate with lack of clinical benefit in ICI-treated NSCLC patients

Focusing on the patient subgroup receiving ICI monotherapy, we identified proteins displaying differential fold changes between R and NR populations. Bioinformatic analysis revealed that the NR dataset was enriched with nuclear and intracellular proteins (FDR q value<0.1 and log2FC>0.1), some potentially originating from alveolar type 1 and type 2 cells (**Figure 4A-B, Figure S6,** and **Table S6**). For example, we found 5 RNA splicing and metabolism-related proteins (PSPC1, KHSRP, EWSR1, SF1, and PUF60), 3 transcriptional regulators (YAP1, HMGA1 and SF1), and 2 metabolite interconversion enzymes (DCPS and ALOX15B) (35) that were elevated in NR compared to R groups. Furthermore, we found that proteins potentially originating from alveolar cells were significantly associated with clinical benefit, but not with other parameters such as age, ECOG, sex, and line of treatment (**Table S7**). Interestingly, patients with high fold change in YAP1, a protein recently shown to be associated with NSCLC and metastasis (36), displayed significantly shorter OS than patients with low fold change in this protein. The survival analysis based on YAP1 yielded the highest HR compared to similar analyses based on other significant proteins expressed by alveolar cells, such as DCPS and DCBLD1 (HR=2.87, p<0.001; **Figure 4C and Figure S7**). Of note, YAP1, TEAD3 and TEAD4 were enriched in the Hippo signaling pathway with the lowest p-value in comparison to intracellular proteins associated with alveolar cells (**Figure 4B** and **Table S7**), further indicating their significant role in NSCLC. To gain further biological insights, we compared fold changes displayed by the R and NR groups within the ICI-treated NSCLC cohort to chemotherapy-treated NSCLC patients and ICI-treated melanoma patients. Interestingly, chemotherapy-treated patients displayed a similar trend to the NR group, while ICI-treated melanoma patients displayed a similar trend to the R group (**Figure 4D** and **Figure S8**). The former finding suggests that proteins elevated in the NR group of ICI-treated NSCLC patients are related to cell stress, a process known to be induced by chemotherapy. The latter finding suggests that the intracellular alveolar-associated proteins are unique to NSCLC NR patients treated with ICIs, as they were not identified in any ICI-treated melanoma patients.

**Figure 4:**
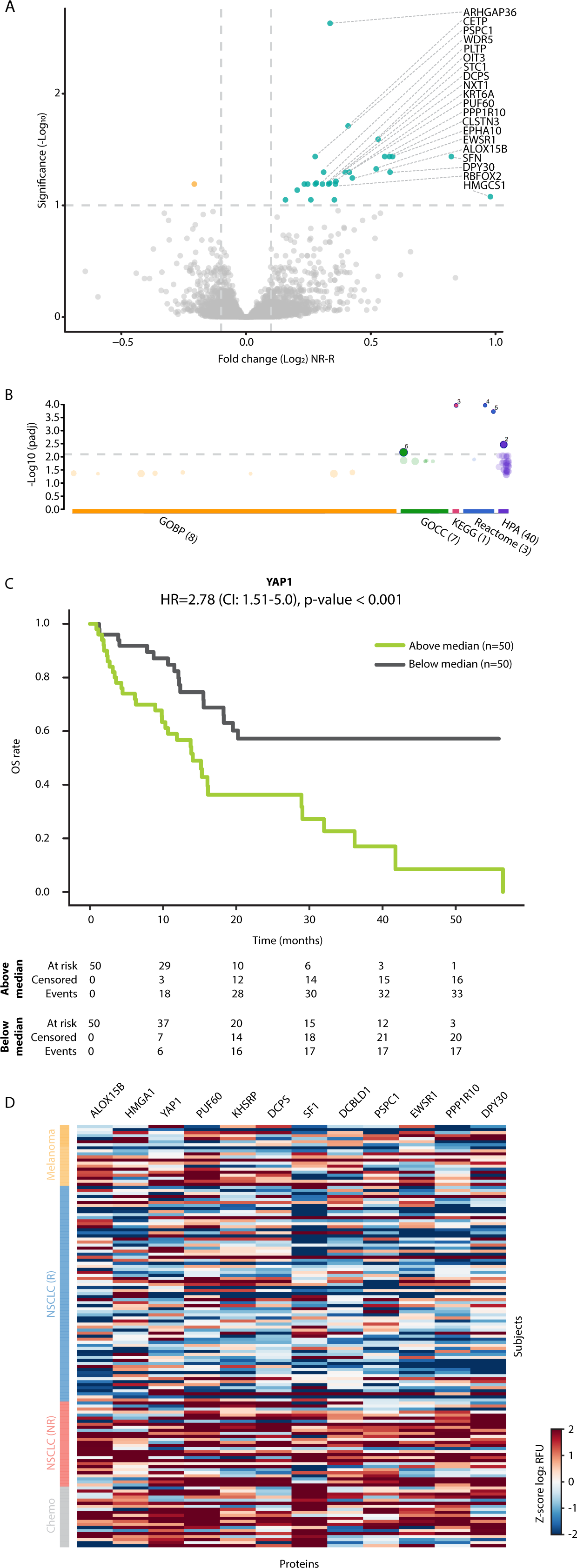
Intracellular alveolar-associated proteins are enriched in plasma of NSCLC patients lacking clinical benefit. A. Volcano plot representing fold change in plasma protein levels [T1:T0 fold change in non-responders (NR) vs T1:T0 fold change in responders (R)] in patients treated with ICI monotherapy. (B) Pathway enrichment analysis of proteins exhibiting significant and positive fold change. Categorization is based on gProfilier. Cut-off was set on - log (q value) of 2.1 (C) Kaplan-Meier plots showing the relationship between YAP1 fold change and overall survival (OS) in patients treated with ICI monotherapy (n=100). Patients were classified as having high or low YAP1 fold change using the median YAP1 fold change as the classification threshold. (D) A heatmap of proteins in R and NR NSCLC patients treated with ICI monotherapy, NSCLC patients treated with chemotherapy and melanoma patients treated with ICI-based therapy. Notably, intracellular alveolar-associated proteins cluster in ICI-treated NR patients and chemotherapy-treated patients.

## DISCUSSION

The current study comprehensively analyzed plasma proteomic profiles in a substantial cohort of 225 NSCLC patients. Leveraging Somalogic aptamer-based technology coupled with bioinformatic approaches, we analyzed approximately 7000 plasma proteins at baseline and during treatment (typically 3-4 weeks following the first treatment dose) to study early on-treatment, systemic proteomic changes. Our study provides insights into the biological mechanisms associated with ICI-based therapies and reveals potential biomarkers for therapeutic benefit. Firstly, we show that the plasma level of sPD-1 is elevated following treatment with PD-1 inhibitors, an effect correlated with better OS, in line with a previous publication (37). sPD-1 elevation is dampened by the addition of chemotherapy to ICI regimens. Secondly, we describe a unique plasma proteomic signature associated with ICI-induced T-cell activation. Thirdly, we identify a group of intracellular proteins potentially expressed by alveolar cells that serve as potential blood-based biomarkers for lack of clinical benefit from ICI therapy.

Focusing on observations related to sPD-1, we show that the fold change in plasma sPD-1 correlates with tumor PD-L1 expression. Specifically, patients with high tumor PD-L1 expression (PD-L1 ≥50%) had a greater fold-change in plasma sPD-1. Interestingly, while sPD-1 fold-change did not correlate with clinical benefit at the 3-month mark, in ICI monotherapy cases, patients with high sPD-1 fold change displayed significantly longer OS than patients with low sPD-1 fold change. These findings are consistent with existing literature reporting that heightened plasma levels of sPD-1 during ICI-based treatment correlate with favorable responses in both EGFR-mutated and wildtype NSCLC patients (37–39). Mechanistically, several possible explanations exist for the increased sPD-1 levels in plasma upon treatment with PD-1 inhibitors. It is plausible that ICI-induced reactivation of cytotoxic T cells augments the proliferation of these cells, thereby fostering increased production of sPD-1 (39–41). However, this explanation does not account for the absence of increased sPD-1 plasma levels upon treatment with PD-L1 inhibitors. In this respect, the therapeutic anti-PD-1 antibodies may bind to sPD-1 in the circulation, interfering with its clearance from the bloodstream, an effect that does not manifest when treatment involves anti-PD-L1 therapy (39). Indeed, the latter explanation aligns with studies on other antibody-based drugs. Specifically, it was documented that upon treatment with anti-IL6 or anti-TNFα, increased circulating levels of IL6 and TNFα, respectively, were detected (42, 43). Overall, our results highlight plasma sPD-1 fold change as a potential predictive indicator for overall survival in patients treated with ICI monotherapy.

The tumor infiltration of T lymphocytes has been previously proposed as a predictive biomarker for ICI therapy outcomes (44). Our study found that upon treatment with ICIs, the plasma proteome is enriched with proteins associated with enhanced T cell activity and proliferation, an effect detectable in both R and NR populations. Conversely, previous studies reported associations between the proliferation of peripheral T lymphocytes or the expression of immune checkpoint molecules such as PD-1 or HAVCR2 (TIM3) and favorable treatment responses (45, 46). Furthermore, a recent study reported increased circulating activated T cells in NSCLC patients upon treatment with immunotherapy (47). Nevertheless, these studies describe the predictive potential of pre-treatment lymphocyte levels in terms of response, progression-free survival or OS (46, 47). Notably, in our study, T cell activation, represented by proteins such as CD8, LAG3, CD28, B2M, GZMA, IL2RA, ILRB, and IL15A, was observed in patients treated with ICI monotherapies, but not combination ICI-chemotherapy. Plausible explanations for this may be related to the immunosuppressive and immune-depleting effects elicited by chemotherapy, which can impact T-cell activity and survival (30, 48, 49). However, this hypothesis does not align with the additive effect seen clinically when ICI and chemotherapy are used concomitantly. An alternative explanation would be that chemotherapy-induced tissue damage within the tumor microenvironment may increase T cell infiltration into the tumor, depleting their presence in the circulation.

An interesting observation from our study is the on-treatment elevation in plasma levels of nuclear and intracellular proteins potentially originating from alveolar cells. Specifically, YAP1 was significantly elevated in NR patients compared to R patients. YAP1 is known to regulate alveolar cell differentiation and induce the Hippo signaling pathway (50). YAP1 is also known to upregulate PD-L1 expression and support immunosuppression (50–52), possibly explaining its upregulation in NR patients in our study. In this regard, our findings suggest a rationale for combining ICIs with a drug that targets YAP1. Importantly, the alveolar-associated proteins measured in the plasma were observed exclusively in the NR population within the cohort of ICI-treated NSCLC patients. Similar proteomic profiles were detected in NSCLC patients treated with chemotherapy alone and were notably absent in ICI-treated melanoma patients. The existence of intracellular, alveolar-associated proteins in the circulation has multiple plausible explanations. Firstly, it is pertinent to acknowledge that larger tumors, often observed in non-responding patients, likely exhibit tumor necrosis and cellular damage, which may impact T-cell activity. Indeed, prior studies have demonstrated that necrotic regions in lung squamous cell carcinoma at baseline may predict an unfavorable response to immunotherapy (53). This can be attributed to the release of intracellular potassium ions from damaged cells that, in turn, affect T cell effector function (54). It is noteworthy, however, that our results specifically reveal on-treatment changes in alveolar proteins, and that such changes are not solely responsible for an attenuated anti-tumor immune response. Secondly, it is possible that the intracellular proteins originate from tumors enduring cellular stress. It is known that chemotherapy exposure induces tumor cell stress, releasing a substantial quantity of extracellular vesicles such as exosomes and microparticles that carry intracellular proteins (55, 56). Therefore, many intracellular proteins described here may originate from extracellular vesicles released upon cell stress. Such proteins will likely be detected in the SomaScan assay, as detergents used in sample preparation may release proteins carried within extracellular vesicles. The finding that such proteins are enriched in the NR population can be explained by cases in which tumor cells remain viable but endure cellular stress, for example when treatment dosage is suboptimal. Alternatively, tumor stroma represented in part by alveolar cells can be tumor protective, and as such, alveolar cells may potentially support lung cancer. However, there is a paucity of data in this area of lung cancer research.

The application of aptamer technology, enabling the profiling of nearly 7000 distinct proteins, holds substantial potential for advancing precision medicine for cancer as well as other diseases. A recent study, using the Somalogic technology identified plasma protein signatures that can predict organ aging in health and disease (24). In contrast to the traditional reliance on cellular or tissue-based biopsies, analyzing plasma proteins through liquid biopsy methodologies offers a more accessible and clinically viable approach. Indeed, our study provides the first comprehensive analysis of plasma proteomes using the Somalogic technology. It successfully identified proteins associated with tumor tissue, including those potentially originating from alveolar cells. Nevertheless, additional technologies are necessary for studying connections between the tumor and circulating proteins. One such avenue involves ctDNA analysis which has the potential to provide valuable insights into tumor-specific features such as genetic mutations and response patterns (57). The integration of distinct technologies, each capturing unique aspects of the tumor and host microenvironment, holds promise for enhancing the predictive power of biomarkers.

## ACKNOWLEDGMENTS

This study was supported by OncoHost. LTD.

## Declaration of competing interests

NR, BR, AA, NN, ALB, ML, RK, AA, MAA, RL, MG, TH, IW, ET, CL, SRS declare no conflict of interest. EJ, GL, BY, YE, IS, ND, MH, CL, YB, MY are employees of Oncohost. JB, APD, YS are consultants of Oncohost

## Authors’ contribution

Conception and design: EJ, GL, BY, YE, IS, YS

Acquisition of data: JB, BR, AA, NN, ALB, ML, RK, AA, MAA, RL, MG, TH, IW, ET, CL, SRS.

Analysis and interpretation of data: EJ, GL, BY, YB, MY, CL, ND, MH, YE, IS, APD, YS.

Writing, reviewing, and revising the manuscript: EJ, GL, ND, MH, APD, YS.

Study supervision: IS, YS

