## Supplemental data for "Biological insights from plasma proteomics of non-small cell lung cancer patients treated with immunotherapy"

**<u>Supplemental data online</u>**

**Supplemental Tables**

**Table S1: Treatment modalities used in the NSCLC patient cohort.**

| Treatment modality | Drugs | Number of patients |
|---|---|---|
| ICI | Pembrolizumab | 69 |
| | Nivolumab | 26 |
| | Atezolizumab | 4 |
| | Durvalumab | 1 |
| ICI + Chemo | Pembrolizumab + Carboplatin + Pemetrexed | 83 |
| | Pembrolizumab + Carboplatin + Paclitaxel | 27 |
| | Pembrolizumab + Cisplatin + Pemetrexed | 9 |
| | Atezolizumab + Bevacizumab + Carboplatin + Paclitaxel | 2 |
| | Pembrolizumab + Nab-Paclitaxel + Carboplatin | 1 |
| | Durvalumab + Carboplatin +Paclitaxel | 1 |
| | Nivolumab, Carboplatin, Paclitaxel | 1 |
| | Nivolumab, Carboplatin, Pemetrexed | 1 |

ICI, PD-1/PD-L1 inhibitors; ICI+Chemo, PD-1/PD-L1 inhibitors combined with chemotherapy.

**Table S2: Chemotherapy-treated NSCLC cohort - demographics, clinicopathological characteristics and outcomes.**

| Metric | Overall |
|---|---|
| **n** | 20 |
| **Age, mean (SD)** | 65.2 (6.4) |
| **ECOG, n (%)** | |
| 0 | 11 (55.0) |
| 1 | 7 (35.0) |
| 2 | 1 (5.0) |
| 3 | 1 (5.0) |
| **Sex, n (%)** | |
| Female | 6 (30.0) |
| Male | 14 (70.0) |
| **Site, n (%)** | |
| Thoraxklinik DE | 20 (100.0) |
| **Line of treatment, n (%)** | |
| Advanced | 16 (80.0) |
| Unknown | 4 (20.0) |
| **Tumor PD-L1 Level, n (%)** | |
| High (≥50%) | 5 (25.0) |
| Low (1%-49%) | 10 (50.0) |
| Negative (<1%) | 5 (25.0) |
| **Histology, n (%)** | |
| Adenocarcinoma | 11 (55.0) |
| Squamous cell lung cancer | 8 (40.0) |
| Other | 1 (5.0) |
| **RECIST criteria, n (%)** | |
| CR, PR, SD | 6 (30.0) |
| PD | 12 (60.0) |
| NA | 2 (10.0) |

CR, complete response; NA, not applicable; PD, progressive disease; PR, partial response; SD, stable disease.

**Table S3: Melanoma cohort - demographics, clinicopathological characteristics and outcomes.**

| Metric | Overall |
|---|---|
| **n** | 20 |
| **Age, mean (SD)** | 65.5 (14.3) |
| **ECOG, n (%)** | |
| 0 | 17 (85.0) |
| 1 | 1 (5.0) |
| Not reported | 2 (10.0) |
| **Sex, n (%)** | |
| Female | 9 (45.0) |
| Male | 11 (55.0) |
| **Site, n (%)** | |
| Hadassa IL | 16 (80.0) |
| Haemek IL | 3 (15.0) |
| Bnai-Zion IL | 1 (5.0) |
| **Line of treatment, n (%)** | |
| First | 13 (65.0) |
| Advanced | 3 (15.0) |
| Unknown | 4 (20.0) |
| **Histology, n (%)** | |
| Cutaneous Melanoma | 10 (50.0) |
| Uveal Melanoma | 4 (20.0) |
| Mucosal Melanoma | 2 (10.0) |
| Acral Melanoma | 1 (5.0) |
| Other/NA | 3 (15.0) |
| **RECIST criteria, n (%)** | |
| CR, PR, SD | 9 (45.0) |
| PD | 9 (45.0) |
| NA | 2 (10.0) |

CR, complete response; NA, not applicable; PD, progressive disease; PR, partial response; SD, stable disease.

**Table S4: Associations between sPD-1 and clinical parameters.**

| Metric | Overall | Low sPD-1 | High sPD-1 | P-value (FDR) |
|---|---|---|---|---|
| **n** | 225 | 113 | 112 | |
| **Age, mean (SD)** | 66.4 (8.2) | 66.0 (8.4) | 66.9 (7.9) | 0.365 (0.657) |
| **ECOG, n (%)** | | | | |
| 0 | 75 (33.3) | 37 (32.7) | 38 (33.9) | 0.548 (0.705) |
| 1 | 128 (56.9) | 64 (56.6) | 64 (57.1) | |
| 2 | 19 (8.4) | 10 (8.8) | 9 (8.0) | |
| 3 | 2 (0.9) | 2 (1.8) | | |
| Not reported | 1 (0.4) | | 1 (0.9) | |
| **Sex, n (%)** | | | | |
| Female | 84 (37.3) | 33 (29.2) | 51 (45.5) | 0.017 (0.051) |
| Male | 141 (62.7) | 80 (70.8) | 61 (54.5) | |
| **Site, n (%)** | | | | |
| Thoraxklinik DE | 98 (43.6) | 40 (35.4) | 58 (51.8) | 0.061 (0.137) |
| Sheba IL | 60 (26.7) | 38 (33.6) | 22 (19.6) | |
| Asklepios DE | 16 (7.1) | 10 (8.8) | 6 (5.4) | |
| Rabin IL | 10 (4.4) | 4 (3.5) | 6 (5.4) | |
| Other | 41 (18.2) | 21 (18.6) | 20 (17.9) | |
| **Line of treatment, n (%)** | | | | |
| First | 165 (73.3) | 84 (74.3) | 81 (72.3) | 0.849 (0.849) |
| Advanced | 60 (26.7) | 29 (25.7) | 31 (27.7) | |
| **Treatment, n (%)** | | | | |
| ICI | 100 (44.4) | 38 (33.6) | 62 (55.4) | 0.002 (0.009) |
| ICI + Chemo | 125 (55.6) | 75 (66.4) | 50 (44.6) | |
| **Tumor PD-L1 Level, n (%)** | | | | |
| High (≥50%) | 89 (39.6) | 32 (28.3) | 57 (50.9) | 0.001 (0.009) |
| Low (1%-49%) | 53 (23.6) | 31 (27.4) | 22 (19.6) | |
| Negative (>1%) | 60 (26.7) | 41 (36.3) | 19 (17.0) | |
| Unknown | 23 (10.2) | 9 (8.0) | 14 (12.5) | |
| **Histology, n (%)** | | | | |
| Adenocarcinoma | 156 (69.3) | 81 (71.7) | 75 (67.0) | 0.519 (0.705) |
| Squamous Cell Carcinoma | 46 (20.4) | 23 (20.4) | 23 (20.5) | |
| Other | 23 (10.2) | 9 (8.0) | 14 (12.5) | |
| **RECIST criteria, n (%)** | | | | |
| CR, PR, SD | 175 (78.4) | 86 (76.7) | 89 (80.1) | 0.650 (0.731) |
| PD | 48 (21.5) | 26 (23.2) | 22 (19.8) | |
| NA | 2 (0.1) | 1 (0.1) | 1 (0.1) | |

NSCLC ICI and ICI+Chemo cohorts were stratified by high and low levels of sPD-1 (9227-15 aptamer). CR, complete response; ICI, PD-1/PD-L1 inhibitors; ICI+Chemo, PD-1/PD-L1 inhibitors combined with chemotherapy; NA, not applicable; PD, progressive disease; PR, partial response; SD, stable disease.

**Table S5: Top biological processes enriched in ICI monotherapy cohort.**

| | Source | Term name | Term id | -log10 (adjusted p value) |
|---|---|---|---|---|
| 1 | GO:BP | immune system process | GO:0002376 | 21.58426 |
| 2 | GO:BP | adaptive immune response | GO:0002250 | 17.88416 |
| 3 | GO:BP | immune response | GO:0006955 | 17.28246 |
| 4 | GO:BP | positive regulation of immune system process | GO:0002684 | 14.00563 |
| 5 | GO:BP | regulation of immune system process | GO:0002682 | 13.29835 |
| 6 | GO:BP | lymphocyte activation | GO:0046649 | 13.10164 |
| 7 | GO:BP | lymphocyte mediated immunity | GO:0002449 | 13.10164 |
| 8 | GO:BP | adaptive immune response based on somatic recombination of immune receptors built from immunoglobulin superfamily domains | GO:0002460 | 12.66585 |
| 9 | GO:BP | cell killing | GO:0001906 | 12.58107 |
| 10 | GO:BP | leukocyte activation | GO:0045321 | 12.35538 |
| 11 | GO:BP | cell activation | GO:0001775 | 12.30476 |
| 12 | GO:BP | T cell activation | GO:0042110 | 11.7129 |
| 13 | GO:BP | leukocyte cell-cell adhesion | GO:0007159 | 11.0257 |
| 14 | GO:BP | regulation of cell killing | GO:0031341 | 10.94615 |
| 15 | GO:BP | leukocyte mediated immunity | GO:0002443 | 10.82761 |
| 16 | GO:BP | immune effector process | GO:0002252 | 10.43794 |
| 17 | GO:BP | regulation of lymphocyte mediated immunity | GO:0002706 | 10.43794 |
| 18 | GO:BP | defense response | GO:0006952 | 10.31962 |
| 19 | GO:CC | external side of plasma membrane | GO:0009897 | 14.23286 |
| 20 | GO:CC | cell surface | GO:0009986 | 14.23286 |
| 21 | GO:CC | plasma membrane | GO:0005886 | 13.42386 |
| 22 | GO:CC | cell periphery | GO:0071944 | 11.91339 |
| 23 | GO:CC | side of membrane | GO:0098552 | 11.77873 |
| 24 | REAC | Immunoregulatory interactions between a Lymphoid and a non-Lymphoid cell | REAC:R-HSA-198933 | 21.89671 |
| 25 | REAC | Immune System | REAC:R-HSA-168256 | 18.69919 |
| 26 | REAC | Adaptive Immune System | REAC:R-HSA-1280218 | 14.75221 |

**Table S6: Top biological processes enriched in non-responders.**

| | Source | Term name | Term id | -log10 (adjusted p value) |
|---|---|---|---|---|
| 1 | HPA | lung; alveolar cells type II | HPA:0301371 | 2.463387355 |
| 2 | HPA | lung; alveolar cells type I | HPA:0301361 | 2.463387355 |
| 3 | KEGG | Hippo signaling pathway - multiple species | KEGG:04392 | 3.959052061 |
| 4 | REAC | RUNX3 regulates YAP1-mediated transcription | REAC:R-HSA-8951671 | 3.964585372 |
| 5 | REAC | YAP1- and WWTR1 (TAZ)-stimulated gene expression | REAC:R-HSA-2032785 | 3.724222096 |
| 6 | GO:CC | nucleus | GO:0005634 | 2.168533572 |

**Table S7: Associations between alveolar proteins and clinical parameters**

| Metric | Overall | Low Alv | High Alv | P-value (FDR) |
|---|---|---|---|---|
| **n** | 100 | 50 | 50 | |
| **Age, mean (SD)** | 67.6 (8.2) | 67.9 (8.0) | 67.2 (8.4) | 0.679 (0.821) |
| **ECOG, n (%)** | | | | |
| 0 | 37 (37.0) | 16 (32.0) | 21 (42.0) | 0.718 (0.821) |
| 1 | 53 (53.0) | 28 (56.0) | 25 (50.0) | |
| 2 | 8 (8.0) | 5 (10.0) | 3 (6.0) | |
| 3 | 2 (2.0) | 1 (2.0) | 1 (2.0) | |
| **Sex, n (%)** | | | | |
| Female | 43 (43.0) | 20 (40.0) | 23 (46.0) | 0.686 (0.821) |
| Male | 57 (57.0) | 30 (60.0) | 27 (54.0) | |
| **Site, n (%)** | | | | |
| Thoraxklinik DE | 48 (48.0) | 26 (52.0) | 22 (44.0) | 0.91 (0.910) |
| Sheba IL | 27 (27.0) | 12 (24.0) | 15 (30.0) | |
| Asklepios DE | 3 (3.0) | 1 (2.0) | 2 (4.0) | |
| Rabin IL | 2 (2.0) | 1 (2.0) | 1 (2.0) | |
| Other | 20 (20.0) | 10 (20.0) | 10 (20.0) | |
| **Line of treatment, n (%)** | | | | |
| First | 53 (53.0) | 29 (58.0) | 24 (48.0) | 0.423 (0.821) |
| Advanced | 47 (47.0) | 21 (42.0) | 26 (52.0) | |
| **Treatment, n (%)** | | | | |
| ICI | 100 (100.0) | 50 (100.0) | 50 (100.0) | |
| **Tumor PD-L1 Level, n (%)** | | | | |
| High (≥50%) | 65 (65.0) | 35 (70.0) | 30 (60.0) | 0.093 (0.372) |
| Low (1%-49%) | 13 (13.0) | 8 (16.0) | 5 (10.0) | |
| Negative (>1%) | 12 (12.0) | 2 (4.0) | 10 (20.0) | |
| Unknown | 10 (10.0) | 5 (10.0) | 5 (10.0) | |
| **Histology, n (%)** | | | | |
| Adenocarcinoma | 68 (68.0) | 32 (64.0) | 36 (72.0) | 0.502 (0.821) |
| Squamous Cell Carcinoma | 25 (25.0) | 15 (30.0) | 10 (20.0) | |
| Other | 7 (7.0) | 3 (6.0) | 4 (8.0) | |
| **RECIST criteria, n (%)** | | | | |
| CR, PR, SD | 71 (71.0) | 46 (92.0) | 25 (50.0) | 1.68E-5 (1.34E-4) |
| PD | 28 (28.0) | 4 (8.0) | 24 (48.0) | |
| NA | 1 (1.0) | 0 (0.0) | 1 (2.0) | |

NSCLC ICI monotherapy cohort stratified by high and low levels of alveolar proteins (median of z-score normalization). Alv, alveolar cells; CR, complete response; ICI, PD-1/PD-L1 inhibitors; NA, not applicable; PD, progressive disease; PR, partial response; SD, stable disease.

### Supplemental Figures

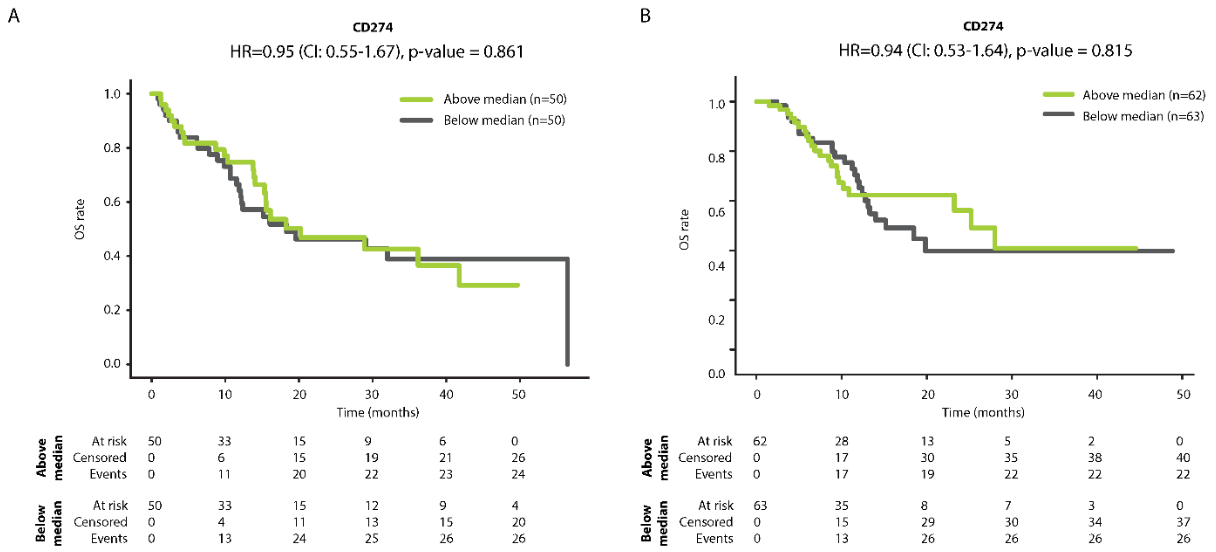


**Figure S1: Kaplan-Meier plots showing the relationship between sPD-L1 fold change and overall survival.** Patients were treated with PD-1/PD-L1 inhibitor monotherapy (n=100; ICI; A) or PD-1/PD-L1 inhibitors in combination with chemotherapy (n=125; ICI+chemo; B). Patients were classified as having high or low sPD-L1 fold change (sPD-L1-high or sPD-L1-low, respectively) using the median sPD-L1 fold change as the classification threshold.

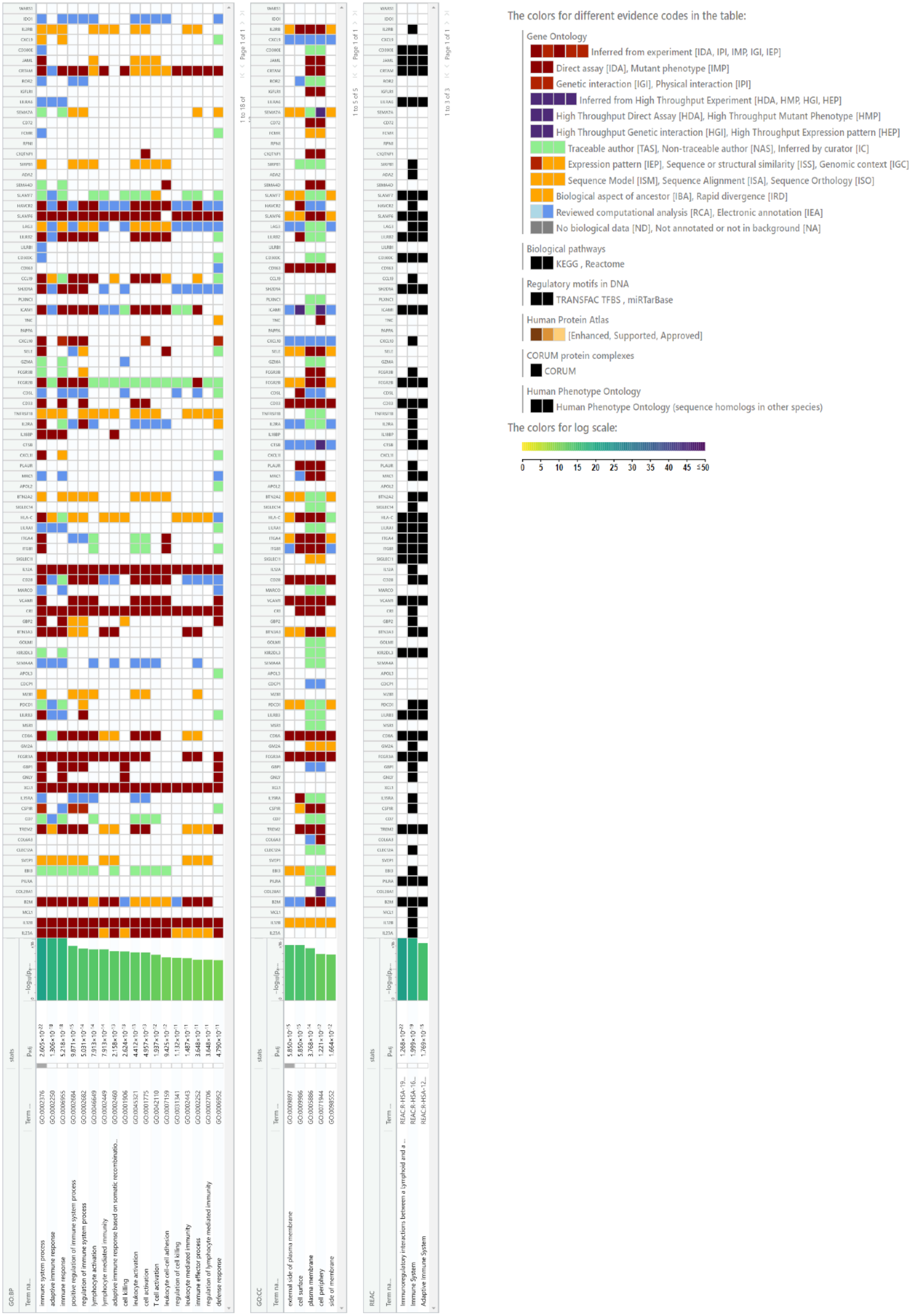
The colors for different evidence codes in the table:
Gene Ontology
Inferred from experiment [IDA, IPI, IMP, IGI, IEP]
Direct assay [IDA], Mutant phenotype [IMP]
Genetic interaction [IGI], Physical interaction [IPI]
Inferred from High Throughput Experiment [HDA, HMP, HGI, HEP]
High Throughput Direct Assay [HDA], High Throughput Mutant Phenotype [HMP]
High Throughput Genetic interaction [HGI], High Throughput Expression pattern [HEP]
Traceable author [TAS], Non-traceable author [NAS], Inferred by curator [IC]
Expression pattern [IEP], Sequence or structural similarity [ISS], Genomic context [IGC]
Sequence Model [ISM], Sequence Alignment [ISA], Sequence Orthology [ISO]
Biological aspect of ancestor [IBA], Rapid divergence [IRD]
Reviewed computational analysis [RCA], Electronic annotation [IEA]
No biological data [ND], Not annotated or not in background [NA]
Biological pathways
KEGG , Reactome
Regulatory motifs in DNA
TRANSFAC TFBS , miRTarBase
Human Protein Atlas
[Enhanced, Supported, Approved]
CORUM protein complexes
CORUM
Human Phenotype Ontology
Human Phenotype Ontology (sequence homologs in other species)
The colors for log scale:
0 5 10 15 20 25 30 35 40 45 ≤50
GO:BP
GO:CC
REAC

**Figure S2: Enrichment of immune-related processes and membrane proteins.** Enrichment analyses were performed on proteins exhibiting significant and positive fold changes (T1:T0) in the ICI monotherapy cohort (n=100). Top GO biological processes and GO cellular compartments are shown.

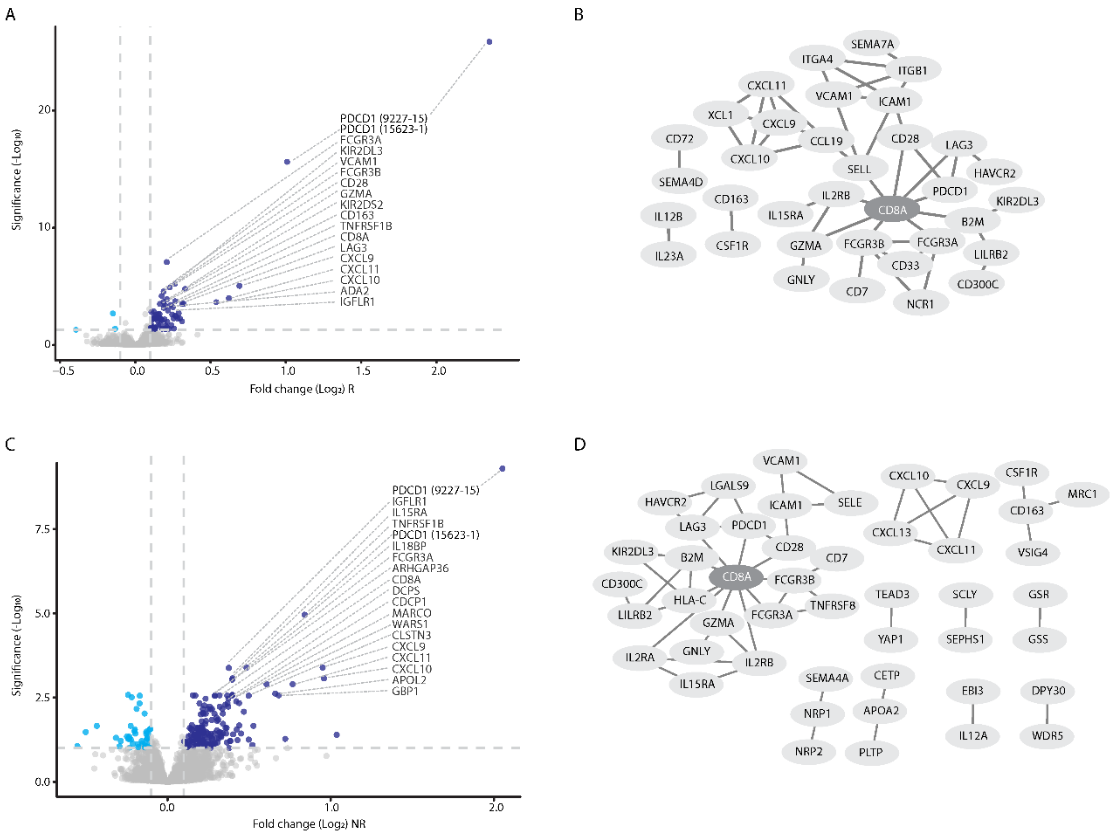


**Figure S3: A T cell hub exists in responder and non-responder NSCLC patients.** Volcano plots representing fold change in plasma protein levels (T1:T0) in patients treated with ICI monotherapy. Responder cohort (n=71; R) is shown in (A) and non-responder cohort (n=28; NR) is shown in (C). Protein-protein interaction networks were generated from proteins with significant and positive fold change in R and NR protein sets, shown in (B) and (D), respectively. Interaction score with 0.9 confidence.

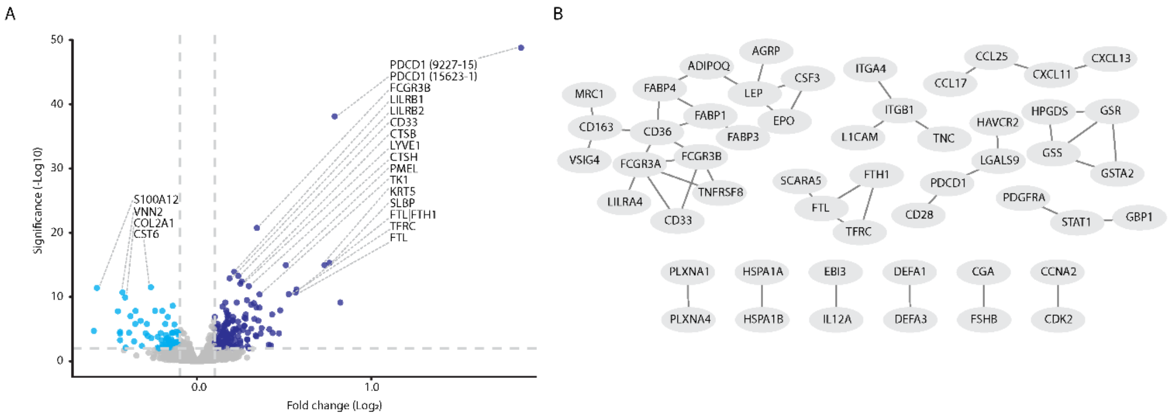


**Figure S4: The addition of chemotherapy to ICIs weakens the T cell hub.** A. Volcano plot representing fold change in plasma protein levels (T1:T0) in patients treated with PD-1/PD-L1 inhibitors combined with chemotherapy (n=125). B. A protein-protein interaction network was generated from proteins with significant and positive fold change. Interaction score with 0.9 confidence. Notably, protein-protein interactions do not present a T cell hub.

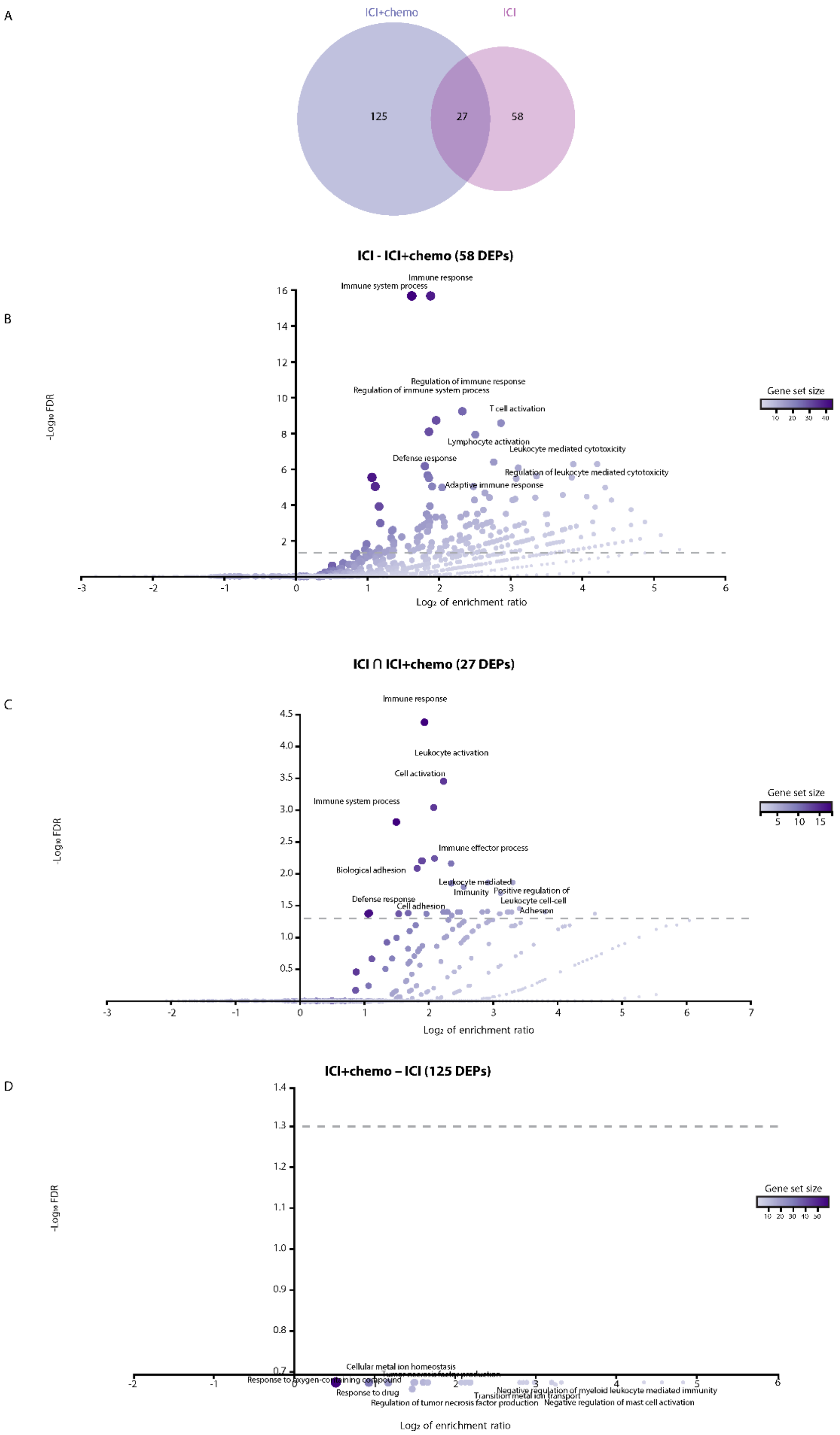
A
ICI+chemo
ICI
125
27
58
B
ICI - ICI+chemo (58 DEPs)
Immune response
Immune system process
Regulation of immune response
Regulation of immune system process
T cell activation
Lymphocyte activation
Leukocyte mediated cytotoxicity
Defense response
Regulation of leukocyte mediated cytotoxicity
Adaptive immune response
Gene set size
-Log10 FDR
Log2 of enrichment ratio
C
ICI ∩ ICI+chemo (27 DEPs)
Immune response
Leukocyte activation
Cell activation
Immune system process
Immune effector process
Biological adhesion
Leukocyte mediated
Immunity
Positive regulation of
Leukocyte cell-cell
Adhesion
Defense response
Cell adhesion
Gene set size
-Log10 FDR
Log2 of enrichment ratio
D
ICI+chemo – ICI (125 DEPs)
Cellular metal ion homeostasis
Response to oxygen-containing compound
Response to drug
Regulation of tumor necrosis factor production
Transition metal ion transport
Negative regulation of myeloid leukocyte mediated immunity
Negative regulation of mast cell activation
Gene set size
-Log10 FDR
Log2 of enrichment ratio

**Figure S5: T cell-related processes are highly enriched in the ICI monotherapy cohort.** A. Venn diagram showing number of proteins with significant and positive fold changes (T1:T0) in ICI monotherapy (ICI) and ICI combined with chemotherapy (ICI+chemo) cohorts. B-D. Pathway enrichment analyses were performed on the following protein sets: 58 proteins exclusive to ICI cohort (B); 27 proteins found in both cohorts (C); 125 proteins exclusive to ICI+chemo cohort (D). The analysis was performed using WebGestalt.

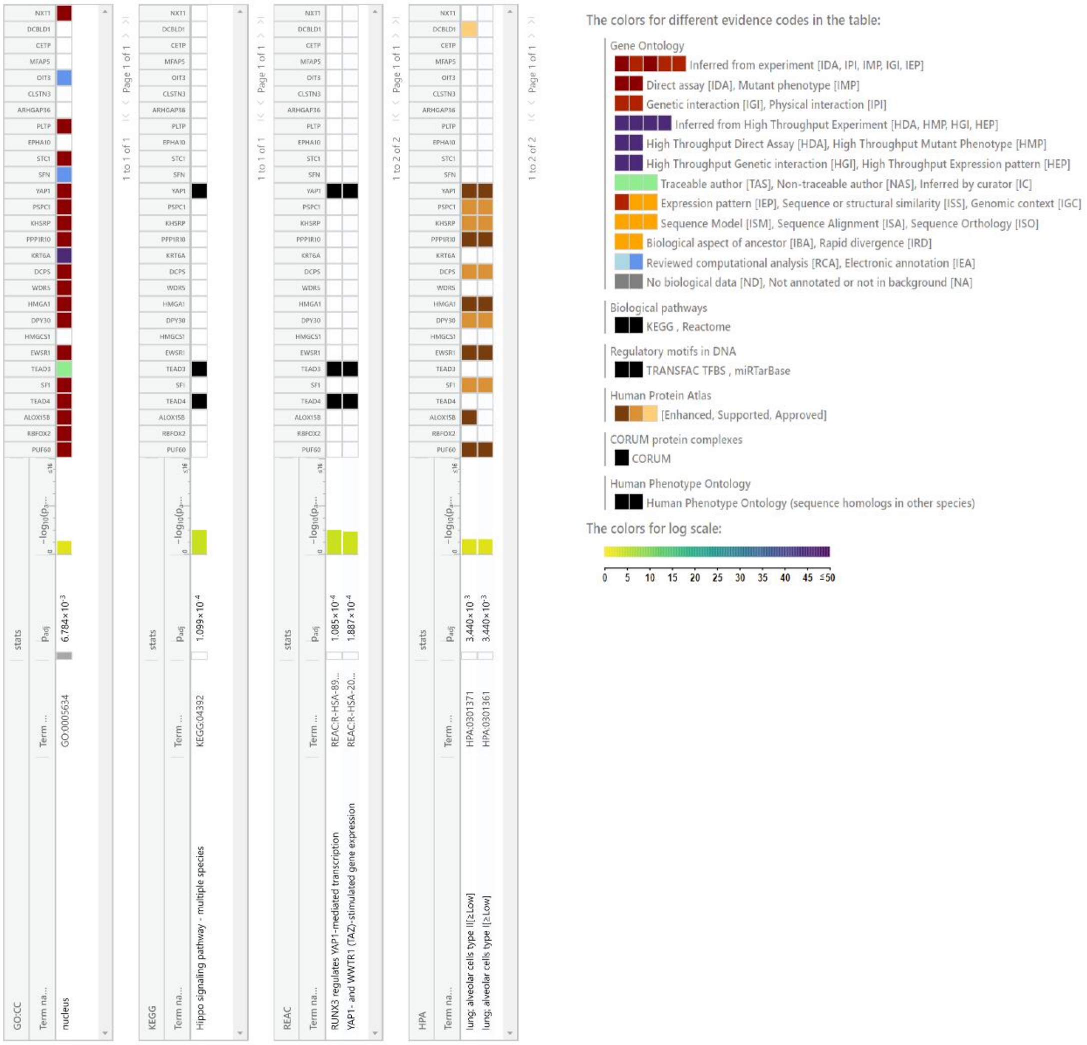


**Figure S6: Nuclear or intracellular proteins are enriched in non-responder NSCLC patients.** Enrichment analyses were performed on proteins exhibiting significant and positive fold changes (T1:T0) in the non-responder subcohort of NSCLC patients treated with ICI monotherapy (n=27). Top results from GO cellular compartments and Human Protein Atlas are shown.

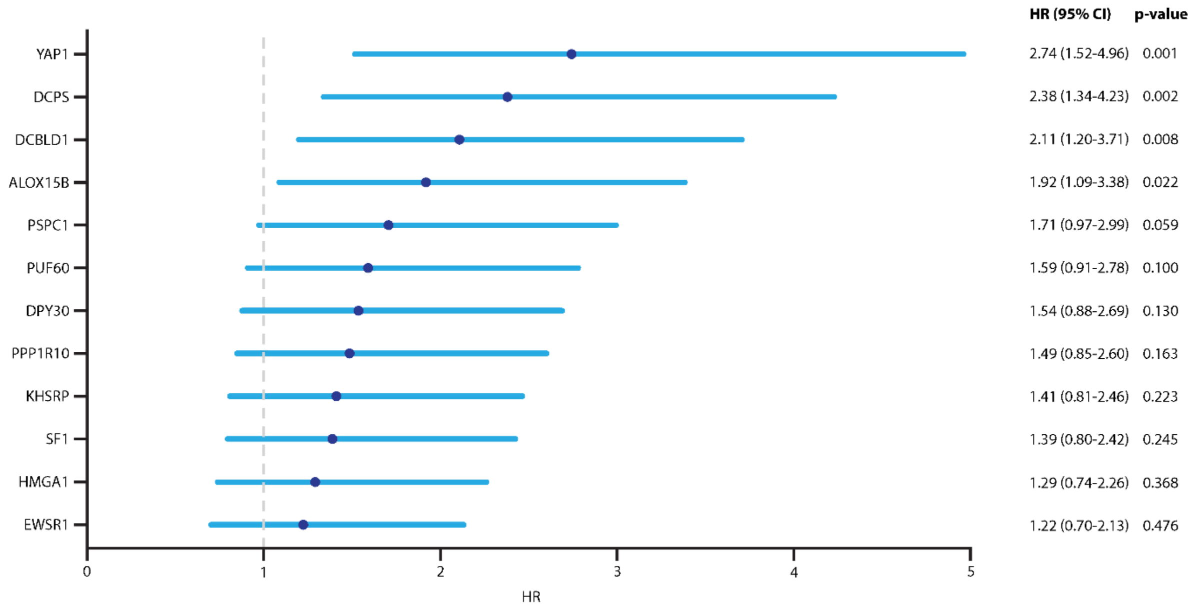


**Figure S7: Kaplan-Meier hazard ratios showing the relationship between selected alveolar proteins and overall survival.** The analysis was performed on the ICI monotherapy patient cohort (n=100). Patients were classified as having high or low fold change for each of the indicated alveolar proteins, using the median fold change as the classification threshold. Hazard ratios (HR) are shown.

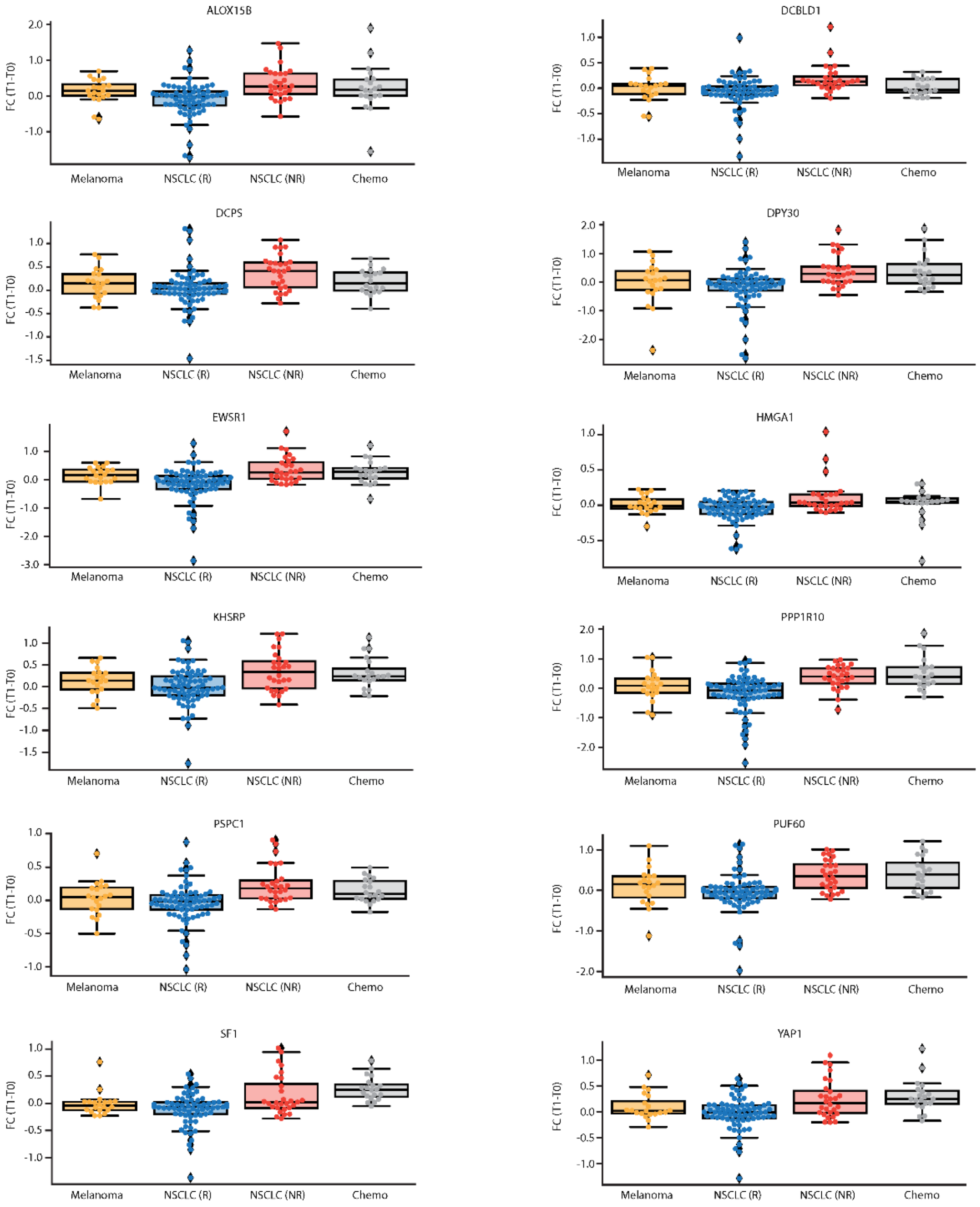


**Figure S8: Fold change of selected alveolar proteins in different patient cohorts.** Proteins (with potential alveolar origin) displaying significantly higher plasma levels in non-responder (NR) compared to responder (R) NSCLC patients treated with ICI monotherapy were selected for analysis in different patient cohorts. Shown are the fold changes (T1:T0) per protein in R and NR NSCLC patients treated with ICI monotherapy, NSCLC patients treated with chemotherapy and melanoma patients treated with ICI-based therapy.
